# Experimental characterization of chicken *OSX/SP7* and embryonic expression analysis reveal skeletal and neural expression domains

**DOI:** 10.64898/2026.08.04.740911

**Authors:** Domink Lonken, Cholpon Zhakshylykova, Clemens Lumper, Jordi Guimera, Rumsha Khan, Max Neukum, Bernhard Hirt, Ulrike Kohler, Christoph Winkler, Andrea Wizenmann

## Abstract

The specificity protein 7 (SP7), also known as Osterix (OSX), is a zinc-finger transcription factor essential for osteoblast differentiation and skeletal development. Hereafter, the protein is referred to as OSX/SP7 throughout the manuscript. Although OSX/SP7 has been extensively studied in mammals and teleost fish, its developmental expression pattern in chicken (*Gallus gallus*) has not been described. Using an experimentally isolated chicken *OSX/SP7* sequence, we examined *OSX/SP7* mRNA expression during embryogenesis by *in situ* hybridization. As expected, *OSX/SP7* expression was detected in developing skeletal elements undergoing ossification. Unexpectedly, transcripts were also observed in the neuroepithelium, retina, central nervous system, embryonic muscles and integument. Notably, *OSX/SP7* expression was present in the neural tube from Hamburger and Hamilton stage 9 and persisted in the developing central nervous system until at least HH40, suggesting that OSX/SP7 functions during avian development may extend beyond osteogenesis. In addition, we reconstructed the chicken *OSX/SP7* coding sequence and inferred its associated untranslated regions. The reconstructed ORF was independently supported by maternal and paternal haplotype-resolved chicken genome assemblies and retained the characteristic domain architecture of vertebrate OSX/SP7 proteins despite substantial divergence outside the DNA-binding domain. Taken together, these findings broaden the developmental landscape of *OSX/SP7* expression in birds and provide new molecular resources for future studies of OSX/SP7 regulation and function during vertebrate development.

## Introduction

The skeletal system is an essential organ system in vertebrates. It provides structural support, protects internal organs, serves as a reservoir of minerals, and together with muscles, enables locomotion. In birds, the skeleton displays several adaptations associated with flight, including pneumatic bones and the fusion of skeletal elements. As in other vertebrates, bone formation occurs through both intramembranous and endochondral ossification (Olsen et al., 2000). Endochondral ossification is particularly important for the development and longitudinal growth of long bones, where a cartilage template is gradually replaced by bone tissue. In chick embryos, this process differs from that of mammals in several aspects, including the absence of secondary ossification centres and a higher degree of vascularization during bone development (Whitehead, 2004).

OSX/SP7 is a zinc-finger-containing transcription factor that plays a central role in osteoblast differentiation and maturation. Studies in mammals have demonstrated that OSX/SP7 is required for bone formation during embryonic development and for bone maintenance in adults (Baek et al., 2010; Huang and Olsen, 2015; Nakashima et al., 2002; Zhou et al., 2010). Similar functions have been reported in teleost fish, including zebrafish and medaka (Niu et al., 2017; Topczewska et al., 2016; Yu et al., 2017). Expression studies in *Xenopus* suggest that this role is evolutionarily conserved among vertebrates (Miura et al., 2008). OSX/SP7 is typically associated with pre-osteoblasts and immature skeletal tissues and is involved in the commitment of mesenchymal progenitors to the osteoblast lineage (Baek et al., 2010; Nakashima et al., 2002; Zhou et al., 2010). The importance of OSX/SP7 for skeletal development is illustrated by loss-of-function studies across different vertebrate species. In humans, mutations in *OSX/SP7* cause an autosomal recessive form of osteogenesis imperfecta characterized by recurrent fractures, bone deformities and delayed tooth eruption (Lapunzina et al., 2010). *OSX/SP7*-deficient mice result in a complete failure of bone formation and fail to form bone despite normal cartilage development (Nakashima et al., 2002), whereas postnatal inactivation results in impaired bone formation and maintenance (Baek et al., 2010). In teleost fish, the severity of *OSX/SP7* mutant phenotypes varies between species, ranging from relatively mild skeletal defects in zebrafish to severe deficiencies in bone formation and juvenile lethality in medaka (Kague et al., 2016; Yu et al., 2017). These observations indicate that, while the osteogenic role of OSX/SP7 is broadly conserved among vertebrates, aspects of its regulation and function may have diverged during evolution.

Although OSX/SP7 is primarily associated with skeletal tissues, its expression is not restricted to bone-forming cells. Expression of *OSX/SP7* has been reported in the olfactory bulb, cortex and cerebellum of mice, in the otic placode of zebrafish, and in otolith-associated tissues of medaka (DeLaurier et al., 2010; Park et al., 2011; Renn and Winkler, 2014). In medaka, OSX/SP7 has been shown to be required for otolith maintenance (Renn and Winkler, 2014), suggesting functions beyond osteoblast differentiation and skeletal development. These findings raise the possibility that OSX/SP7 may participate in developmental processes outside the skeletal system.

Predicted *OSX/SP7* -related sequences are present in the chicken genome, but experimental characterization of avian *OSX/SP7* and its embryonic expression pattern has remained limited. In the present study, we experimentally characterized a chicken *OSX/SP7*-related cDNA sequence and used it to generate probes for mRNA in situ hybridization. We analysed the spatial expression pattern of *OSX/SP7* during embryonic development and identified expression domains in both skeletal and non-skeletal tissues. These data provide a foundation for future studies of OSX/SP7 function during avian development.

## Methods

### *Gallus gallus* embryos and tissue sections

Chicken eggs were obtained from LSL Rhein Main and incubated at 38°C and 55% humidity until the required Hamburger-Hamilton (HH) developmental stages were reached (Hamburger and Hamilton, 1951). Eggs were opened at embryonic day (E) 1, E5, E9 and E12 and embryos were isolated and fixed with 4% paraformaldehyde (PFA). Older embryos were embedded in Tissue-Tek OCT compound (Labtech) and cut into 40 µm transverse sections.

### Isolation of a partial chicken OSX/SP7 cDNA

At various stages of embryonic development, total RNA was isolated from embryonic upper leg tissue, including the developing femur, using the RNeasy Mini Kit (Qiagen). First-strand cDNA was synthesized using Maxima H Minus Reverse Transcriptase (Thermo Fisher Scientific) in the presence of RiboLock™ RNase inhibitor (Thermo Fisher Scientific).

Two sets of primers were designed based on the predicted chicken OSX/SP7 sequence (XM_015300329.2) and synthesized by Eurofins Genomics (Germany) for nested PCR amplification.

Primary PCR primers: forward1: 5′-GCGGCGGCGGTGCTGCTCC-3′; reverse1: 5′-GAGCGCGGGCTGAGCCTGC-3′; Nested PCR primers: forward2: 5′-GCTGCTCCGGCAGCGGACC-3′; reverse2: 5′-GGGTGCACGTCCCACCAAGC-3’.

Forward and reverse primers were positioned in different exons to minimize amplification from contaminating genomic DNA and to increase specificity for *OSX/SP7* transcripts. Because of the high GC content of the target region, PCR reactions were performed in the presence of 1.5% DMSO using annealing temperatures of 67°C (Frey et al., 2008; Jensen et al., 2010; Mamedov et al., 2008; Yourno, 1996; Zhang et al., 2009). The primary PCR generated a 611bp fragment, which was purified from agarose gels using the GeneJET Gel Extraction Kit (Thermo Fisher Scientific) and subsequently used as template for nested PCR amplification. The nested PCR yielded a 527 bp fragment (Figure 1), which was gel-purified, cloned into pBluescript II KS+, and sequenced by Sanger sequencing (GATC Biotech, Germany).

**Figure 1.**
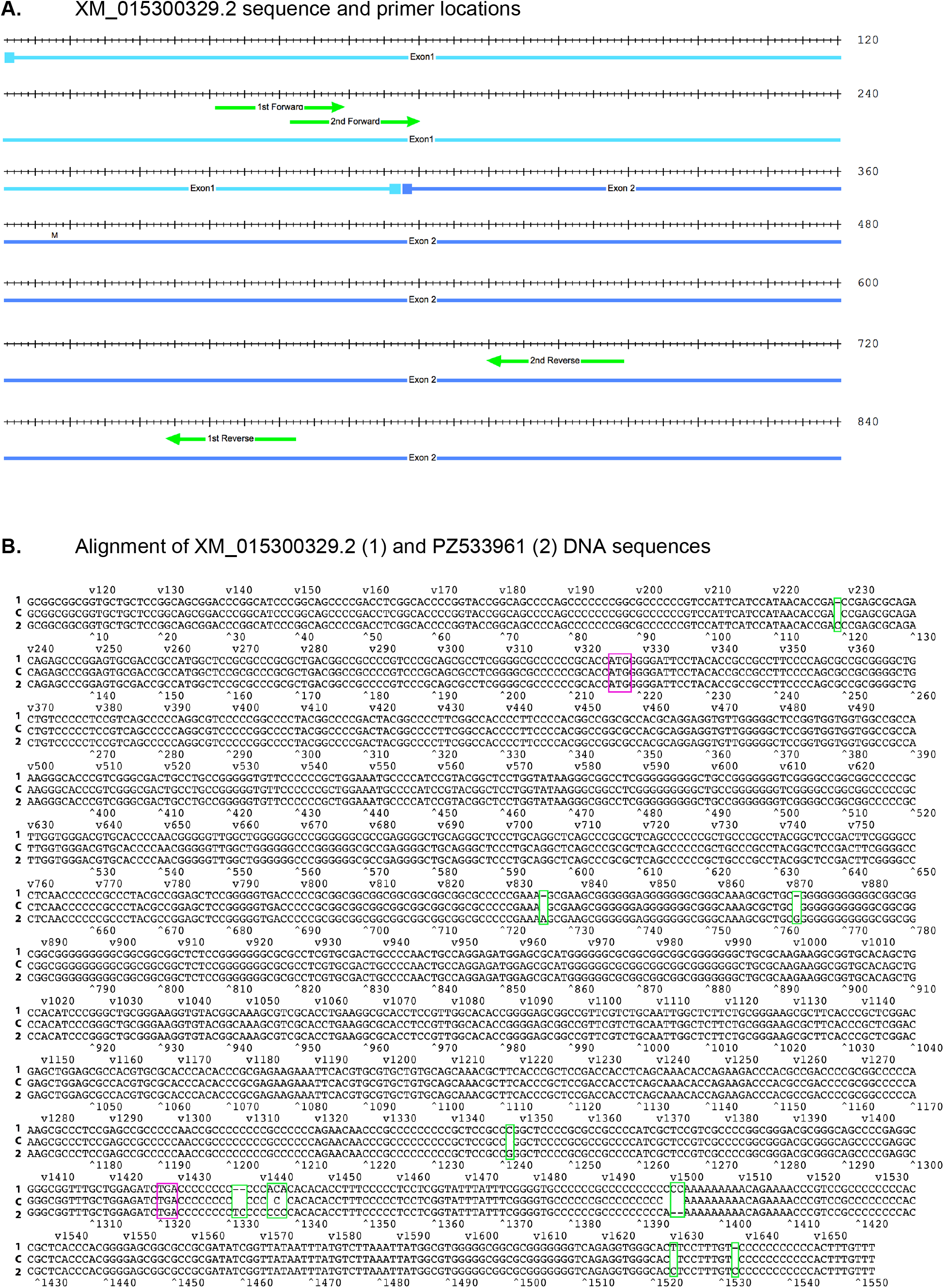
*Gallus gallus OSX/SP7* DNA sequences. **A**: The isolated *OSX/SP7* cDNA fragment is 527 bp long and was obtained by nested PCR. The XM_015300329.2 was used to create the forward and reverse primers. The positions of the primary and nested PCR primers are indicated. The fragment contains the translation initiation codon (ATG) of the predicted chicken OSX/SP7 protein. **B**: The isolated *OSX/SP7* DNA was confirmed by Sanger sequencing and deposited in GenBank under accession number PZ533961. Alignment with the predicted chicken *OSX/SP7* sequence (XM_015300329.2) revealed almost complete identity, with 3 single nucleotide gaps (indicated with green quarters) between the PZ533961 (1 – 1552bp) and XM_015300329.2 (111 – 1658bp) sequences. Start (ATG) and stop (TGA) codon are indicated with violet quarters.

### RNA probe synthesis

Digoxigenin-labelled antisense RNA probes were generated *by in vitro* transcription from the cloned *OSX/SP7* cDNA fragment and used for *in situ* hybridization experiments. The RNA probe was synthesized according to the MBI Fermentas Protocol by RNA polymerase (T7, Invitrogen) by *in vitro* transcription.

### *In situ* hybridisation

*In situ* hybridization was performed essentially as described by Li et al. (Li et al., 2005). Whole-mount *in situ* hybridization was carried out on E1.5 chick embryos, whereas section *in situ* hybridization was performed on 40 µm cryosections obtained from E5, E9 and E12 embryos. Digoxigenin-labelled antisense RNA probes were detected with a peroxidase-coupled DIG antibody (Roche Diagnostics GmbH, Mannheim). BCIP/NBT or Fast Red (Roche Diagnostics GmbH, Mannheim) were used as substrates for the staining reaction catalysed by peroxidase. As a positive control for the hybridization procedure, an *ENGRAILED-2 (EN2*) probe (Logan et al., 1996) was used.

## Results

### Isolation of partial *OSX/SP7* sequence from cDNA

A predicted *OSX/SP7* sequence is present in the chicken genome and is annotated in the NCBI database under accession XM_015300329.2. This sequence was used to design primers for the isolation of a partial *OSX/SP7* cDNA fragment from embryonic cDNA by nested PCR (Fig. 1A).

The experimentally isolated *OSX/SP7* fragment was subsequently used to generate probes for expression analysis. The amplified fragment comprised 526 bp spanning the first two exons of the predicted *OSX/SP7* transcript and included the predicted translation initiation codon. Following cloning, Sanger sequencing confirmed the identity of the amplified fragment. Alignment with the predicted chicken *OSX/SP7* sequence (XM_015300329.2) revealed near-complete sequence identity, with 525 of 526 nucleotides matching the predicted sequence (Fig. 1B). The chicken *OSX/SP7* cDNA sequence generated in this study has been deposited in GenBank under accession number PZ533961.

### Structural and comparative analysis of chicken OSX/SP7

Comparison of the experimentally isolated *OSX/SP7* sequence with recently available genomic resources enabled reconstruction of the complete chicken *OSX/SP7* coding sequence and the inference of its associated untranslated regions from genomic data. Importantly, the same coding sequence was identified independently in both maternal and paternal haplotype-resolved chicken genome assemblies, providing strong support for the reconstructed ORF.

The reconstructed chicken *OSX/SP7* coding sequence encodes a 368-amino acid protein that retains the characteristic domain organization of vertebrate OSX/SP7 proteins despite substantial differences in overall length among species. Human OSX/SP7 comprises 431 amino acids, whereas the murine orthologue encodes a 428-amino acid protein (Nakashima et al., 2002). Predicted OSX/SP7 proteins from other avian species appear shorter, comprising approximately 330 amino acids in species such as *Parus major* and *Lonchura striata*.

Despite these differences in protein length and the divergence observed at the primary sequence level, the reconstructed chicken OSX/SP7 protein retained all major structural features characteristic of vertebrate OSX/SP7 proteins. These included an N-terminal proline-rich region implicated in transcriptional activation, the highly conserved BTD (buttonhead) box located immediately upstream of the DNA-binding domain, the three C2H2 zinc fingers responsible for DNA binding and regulation of osteogenic target genes, and the conserved C-terminal motif GSPEAGGLLEI, which has been associated with structural stability of the zinc-finger domain and interactions with transcriptional cofactors. Taken together, the reconstructed chicken *OSX/SP7* sequence preserves the canonical domain architecture of OSX/SP7 proteins despite substantial sequence divergence outside the DNA-binding domain.

Comparative sequence analysis revealed a high degree of conservation among avian *OSX/SP7* sequences. Alignments of the isolated chicken OSX/SP7 amino-acid sequence with OSX/SP7 sequences from other birds, including *Lonchura striata, Parus major, Numida meleagris* and *Cuculus canorus*, showed sequence similarities of up to 94% (Fig. 2A). Phylogenetic analysis based on the corresponding coding sequences grouped chicken *OSX/SP7* with other avian *OSX/SP7* sequences (Fig. 2B).

**Figure 2.**
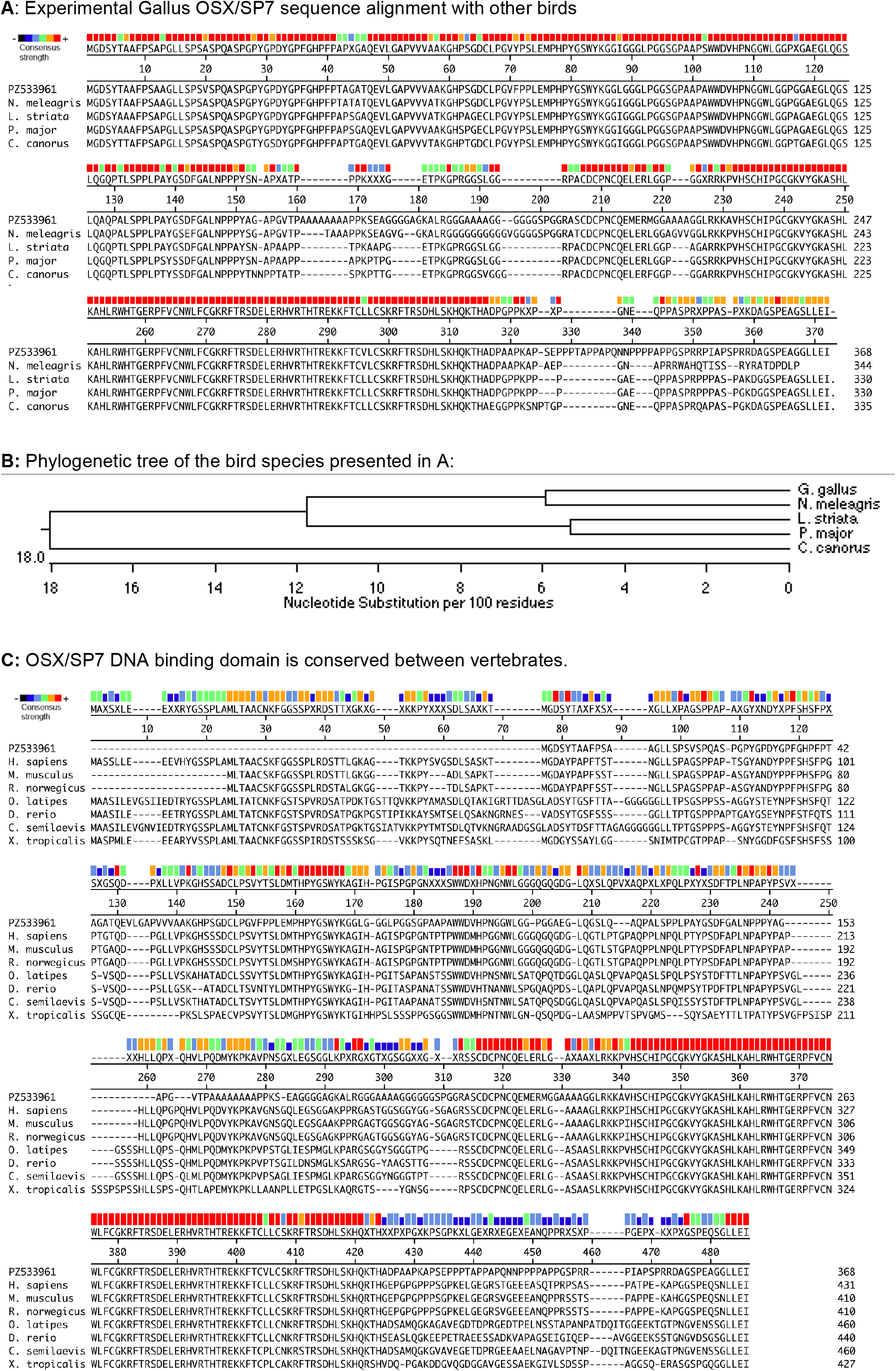
Sequence comparison and phylogenetic analysis of chicken OSX/SP7. **A:** Amino acid sequence alignment of the isolated chicken OSX/SP7 (PZ533961) with OSX/SP7 proteins from different bird species. Two highly conserved regions are highlighted (red boxes above the alignment). The OSX/SP7 sequence of *Numida meleagris* shows the highest overall similarity to chicken OSX/SP7, with sequence conservation extending beyond the two highly conserved regions. **B**: Phylogenetic tree based on nucleotide sequences corresponding to the coding region of avian *OSX/SP7* cDNAs. Branch lengths represent nucleotide substitutions. Among the avian species analysed, the *OSX/SP7* sequence of *Cuculus canorus* is the most divergent from that of *Gallus gallus*. **C**: Amino-acid sequence alignment of chicken OSX/SP7 with its vertebrate homologues from mammals, fish and other vertebrates. Although the overall sequence similarity is lower than that observed among birds, the conserved C-terminal region identified in avian OSX/SP7 proteins is also present in other vertebrate groups. Based on conserved-domain analysis using NCBI Conserved Domain BLAST (https://www.ncbi.nlm.nih.gov/Structure/cdd/wrpsb.cgi; accessed July 2025), this region is predicted to mediate DNA binding. **Abbreviations:** *C. canorus, Cuculus canorus; C. semilaevis, Cynoglossus semilaevis; D. rerio, Danio rerio; G. gallus, Gallus gallus; H. sapiens, Homo sapiens; L. striata, Lonchura striata; M. musculus, Mus musculus; N. meleagris, Numida meleagris; O. latipes, Oryzias latipes; P. major, Parus major; R. norvegicus, Rattus norvegicus; X. tropicalis, Xenopus tropicali*s.

Comparison with OSX/SP7 sequences from mammals (*Mus musculus, Rattus norvegicus* and *Homo sapiens*) and other vertebrates, including *Danio rerio, Oryzias latipes* and *Xenopus tropicalis*, revealed substantially lower levels of amino acid sequence similarity (Fig. 2C). Sequence identity was approximately 55% with teleost fish orthologues and approximately 40% with mammalian orthologues. The highest degree of conservation was found within the C-terminal region of the chicken OSX/SP7 protein. Conserved domain analysis identified this region as the zinc-finger DNA-binding domain characteristic of OSX/SP7 transcription factors. In addition to the highly conserved zinc-finger DNA-binding domain, the reconstructed chicken OSX/SP7 protein retained the characteristic domain organization of vertebrate OSX/SP7 proteins, including an N-terminal proline-rich region, the BTD box, and the conserved C-terminal GSPEAGGLLEI motif.

Taken together, the comparative alignment of OSX/SP7 amino acid sequences revealed a high degree of conservation among vertebrates. Mammalian OSX/SP7 proteins shared 95-99% amino acid identity, whereas avian sequences exceeded 88% identity. In contrast, the chicken OSX/SP7 protein displayed approximately 40% identity with mammalian orthologues.

### Early embryonic expression of OSX/SP7

To determine the onset of *OSX/SP7* expression during embryonic development, *whole-mount* mRNA *in situ* hybridization was performed on chick embryos between HH9 and HH12 (Fig. 3).

**Figure 3.**
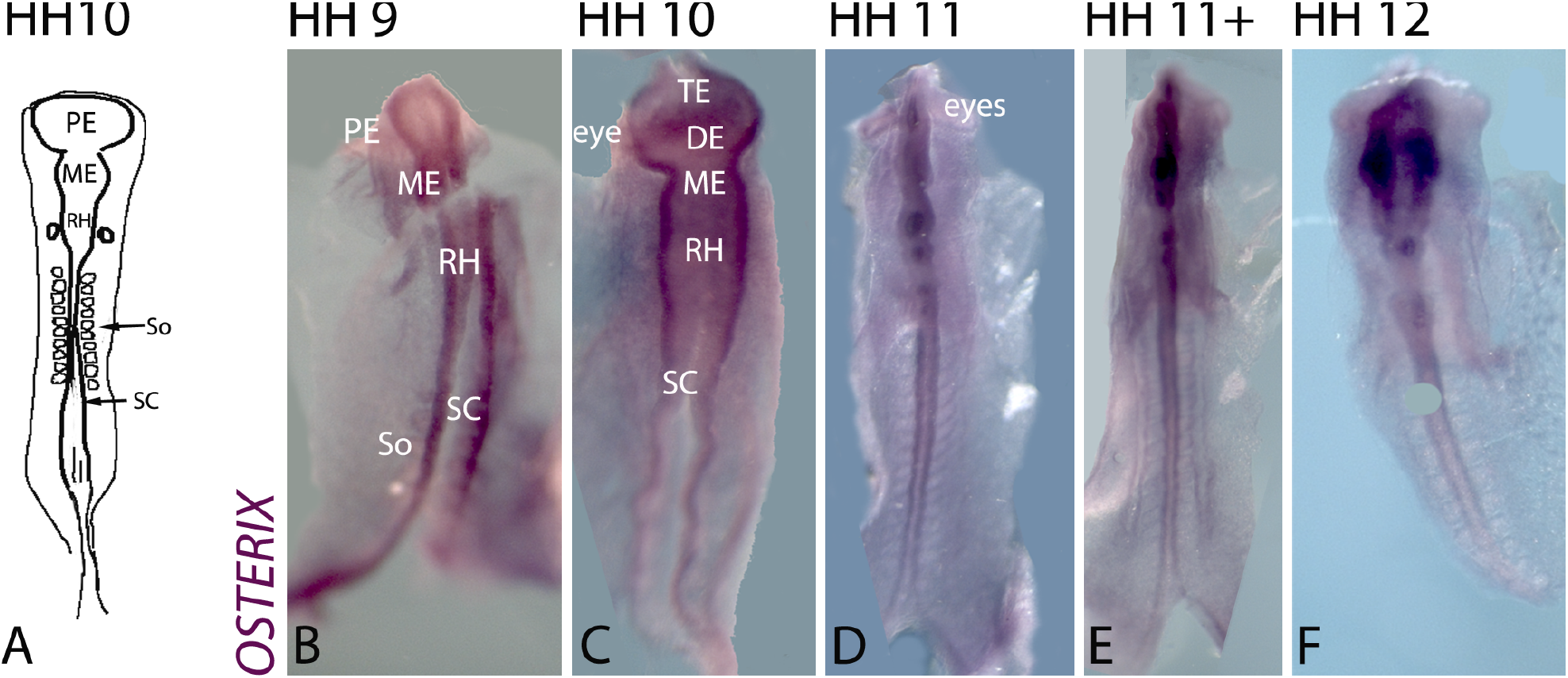
Whole-mount *in situ* hybridization showing *OSX/SP7* expression in early chick embryos. **A**: Schematic drawing of an HH10 embryo indicating the major regions of the developing central nervous system. The different brain regions are indicated. **B-F**: Whole-mount *in situ* hybridization of chick embryos between embryonic day 1.5 and E2 (HH9-HH12). At all stages examined, *OSX/SP7* is expressed in the neuroepithelium of the neural tube along the entire anterior-posterior axis, including all major brain regions and the spinal cord. Weak *OSX/SP7* expression is also visible at the boundaries between adjacent somites at HH9-HH11 (arrow in B). **Abbreviations:** DE, diencephalon; ME, mesencephalon; PE, prosencephalon; RH, rhombencephalon; SC, spinal cord; So, somites. Scale bar: 100 µm.

*OSX/SP7* mRNA expression was first detected at HH9 (E1.5) in the neuroepithelium of the developing neural tube. Expression extended along the entire anterior-posterior axis of the central nervous system, including the prosencephalon, diencephalon, mesencephalon, rhombencephalon and spinal cord (Fig. 3A, B). At HH10, HH11 and HH12, *OSX/SP7* expression remained detectable throughout the developing neural tube and brain regions (Fig. 3C-F). Although regional differences in signal intensity were observed, expression persisted throughout these stages within the developing central nervous system.

In addition to the neural tube, weak *OSX/SP7* expression was observed at somite boundaries between HH9 and HH11 (Fig. 3B-E, arrow). No detectable expression was observed in the developing optic vesicles (Fig. 3C-E, arrowheads).

These observations indicate that *OSX/SP7* expression is initiated at very early stages of chick embryogenesis and is predominantly associated with developing neural tissues.

### Expression of *OSX/SP7* in neural, skeletal, muscular and epithelial tissues during later embryonic development

To examine *OSX/SP7* expression at later stages of embryonic development, mRNA *in situ* hybridization was performed on tissue sections from HH28 to HH40 embryos (Figs. 4-6).

**Figure 4.**
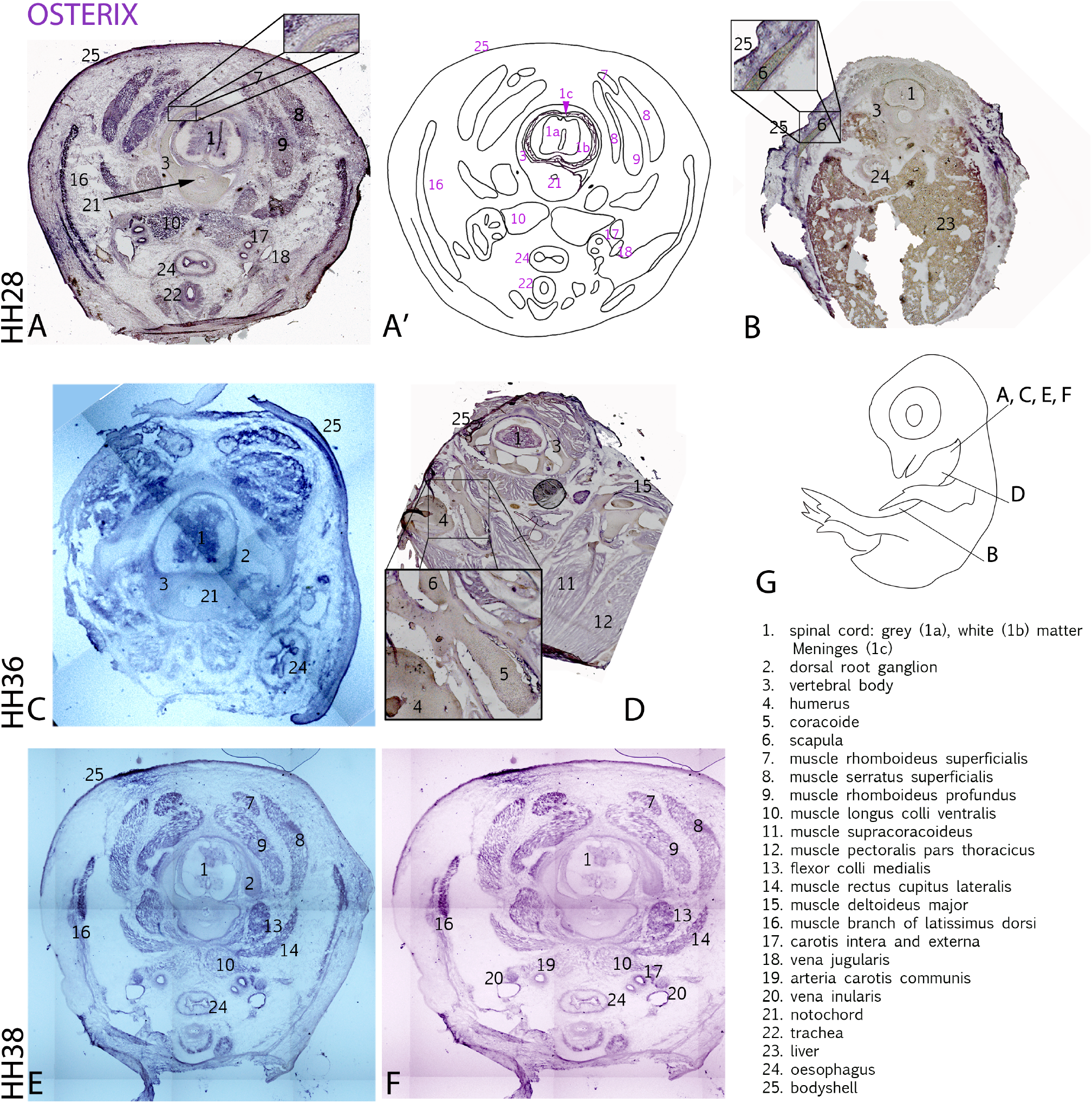
*OSX/SP7* expression in the trunk and shoulder region of chick embryos. **A-F** Transverse sections showing *OSX/SP7* expression at HH28 (E5), HH36 (E10) and HH38 (E12). **G** Schematic drawing of an E11 chick embryo indicating the approximate positions of the transverse sections (40 µm) along the anterior-posterior body axis. Numbered structures are identified in the figure. *OSX/SP7* expression is detected in the spinal cord (1), several muscle groups (7-12, 15, 16), and epithelial tissues including the trachea (22), oesophagus (24), and body wall/integument (25). Expression is also present in skeletal tissues associated with the developing vertebral body (3; magnified in A), on the outside of the scapula (6; magnified in B and D), coracoid (5), and humerus (4; magnified in D). At HH36 and HH38, strong *OSX/SP7* expression is maintained in the spinal cord, muscles and oesophageal epithelium. Cells surrounding the neural tissue of the spinal cord show little or no detectable *OSX/SP7* expression.

Tissue sections from HH28 (E5) to HH38 (E12) showed strong *OSX/SP7* mRNA expression in muscle tissues (Fig. 4A, C, E, F). Expression was visible in the rhomboideus superficialis (label 7), rhomboideus profundus (label 9), serratus superficialis (label 8), deltoideus major (label 15) and in a branch of the latissimus dorsi muscle (label 16). Expression was also detected in neural tissue of the spinal cord (label 1), in the epithelium of the oesophagus (label 24) and trachea (label 22), and in the vena jugularis (label 18). Cells surrounding the neural tissue of the spinal cord showed little or no detectable *OSX/SP7* expression.

Expression associated with skeletal tissues was observed around the vertebral body at HH28, HH36 and HH38 (for example, see the magnification in Fig. 4A) and around the scapula (see magnifications in Fig. 4B and D). At HH36 (E10), the spinal cord, muscles and oesophageal epithelium still showed strong *OSX/SP7* expression. At this stage (Fig. 4D), expression was also visible around the coracoid (label 5) and the humerus (label 4). Strong expression in muscles, epithelia and the spinal cord was still evident at HH38.

Expression studies in brain sections showed that *OSX/SP7* was also strongly expressed in neural tissues of the brain (Fig. 5). At HH34 (E8), both the outer, more differentiated layers and the inner proliferative layers of the optic lobe displayed strong *OSX/SP7* expression (label 1, Fig. 5A). Expression was also detected in the postorbital cartilage and skull bones (label 8 and 21).

**Figure 5.**
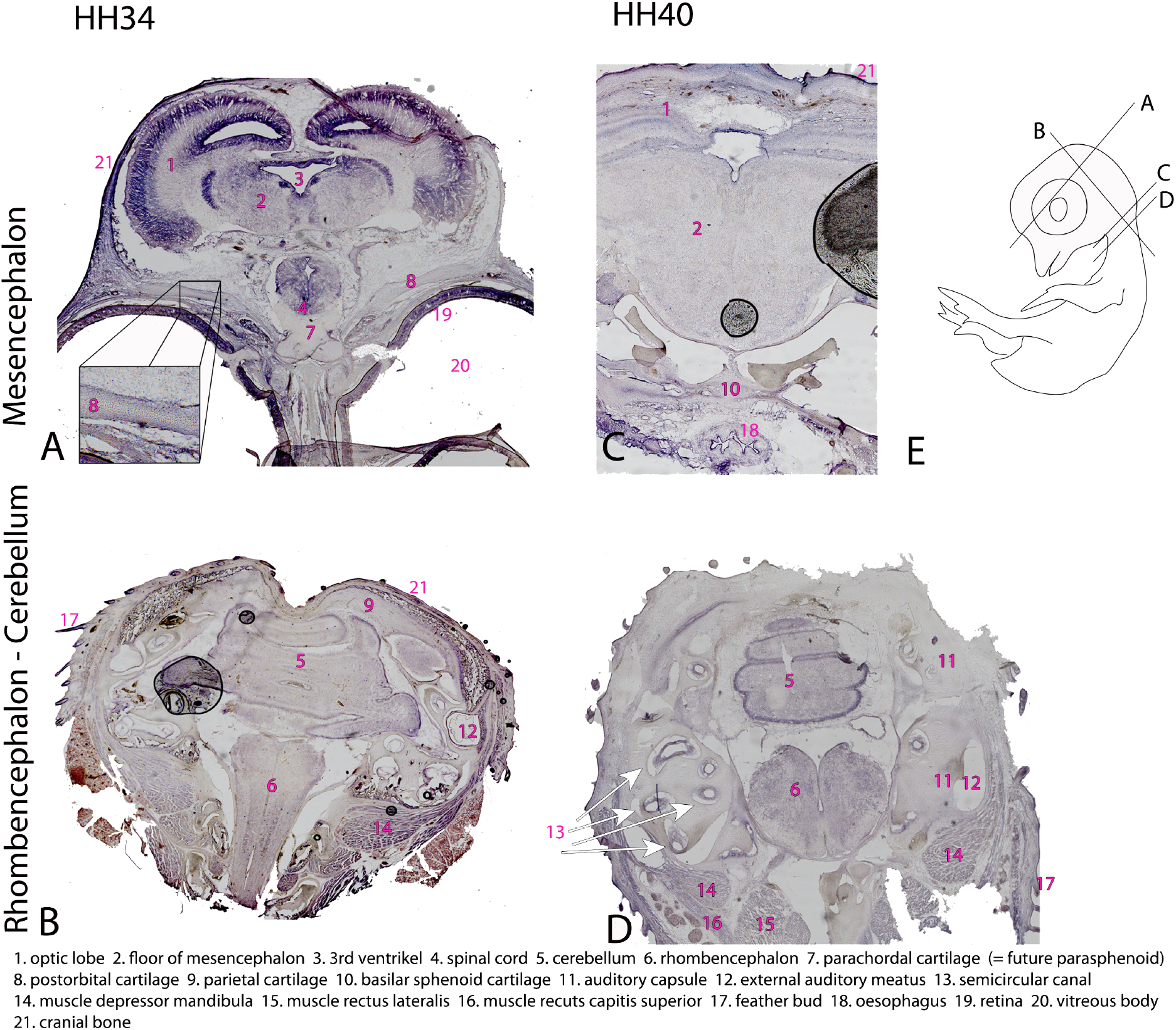
*OSX/SP7* expression in the developing head. **A-B** Sections through the head at HH34 (E8). **C-D** Corresponding sections through the head at HH40 (E15). **E** Schematic drawing of an embryo indicating the approximate position and orientation of the sections shown in A-D. Numbered structures are identified in the figure. At HH34, strong *OSX/SP7* expression is observed in the outer layers of the mesencephalon (1), in neuronal nuclei (2), the spinal cord (4), the retina (19), the postorbital cartilage (8), and cranial bone tissue (21). Weaker expression is present in neurons of the rhombencephalon (6) and in the epithelium surrounding the cerebellum (5). Expression is also detected in the depressor mandibulae muscle (14), feather buds (17), and around the external auditory meatus (12). At HH40 (C), *OSX/SP7* expression persists in several layers of the mesencephalon (1), around the basisphenoid cartilage (10), and in the oesophageal epithelium (18). Expression in ventral mesencephalic neurons (2) is weaker than at HH34. Strong expression remains in the epithelium surrounding the cerebellum (5 in D), whereas several neuronal populations within the rhombencephalon (6) continue to express *OSX/SP7*. Expression is also detected in muscle tissues (14-16) and in the semicircular canal of the inner ear (13).

More posteriorly, in the cerebellum and hindbrain, *OSX/SP7* expression was weak or barely detectable (Fig. 5B). Strong expression was still observed in several muscles, including the depressor mandibulae muscle (label 14), as well as in feather buds (label 17) and around the external auditory meatus (label 12). Six stages later (approximately three days), at HH40, some layers of the optic lobe still showed weak *OSX/SP7* expression, as did the oesophageal epithelium, the basisphenoid cartilage (label 10) and developing skull tissues (Fig. 5C). At HH40, *OSX/SP7* expression was still present in the epithelium surrounding the cerebellum (label 5, Fig. 5D). Weak expression could also be observed in several muscles and feather buds (Fig. 5D). Strong *OSX/SP7* expression was detected in the semicircular canal of the inner ear (label 13, Fig. 5D).

In forelimb sections at HH35 (Fig. 6), strong *OSX/SP7* expression was observed around the developing limb bones. *OSX/SP7* expression was present in the periosteum of the ulna, radius and humerus (label1, 2 and 3). It was also detected in the integument (label 8), whereas expression in muscles such as the extensor metacarpi ulnaris and flexor carpi ulnaris was weak.

**Figure 6.**
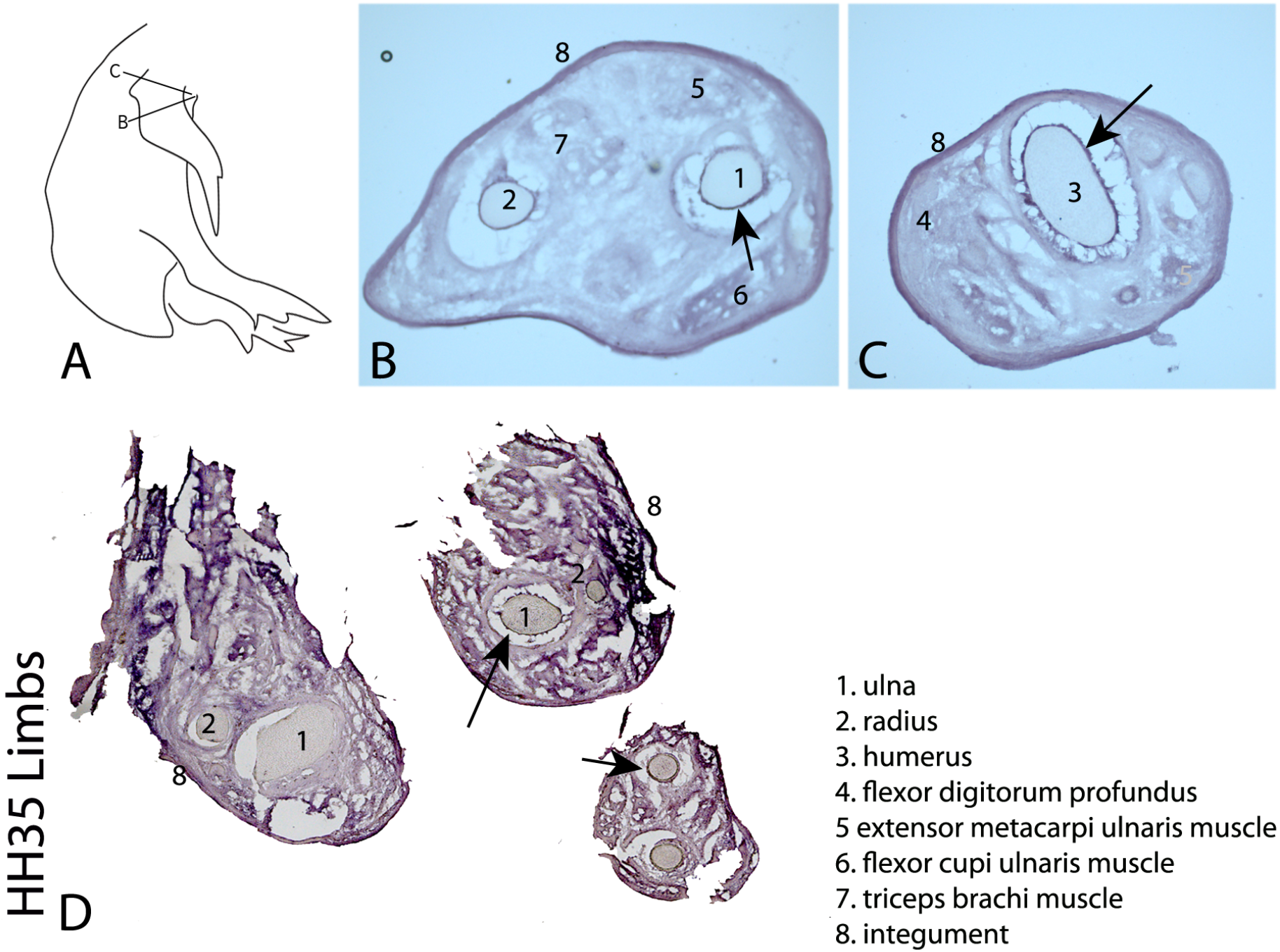
*OSX/SP7* expression in the developing forelimb at HH35 (E9). **A** Schematic drawing indicating the approximate position of the sections shown in B-C. Numbered structures are identified in the figure. **B-D** Transverse sections through the forelimb. Strong *OSX/SP7* expression is detected around the ulna (1), radius (2), and humerus (3) (arrows). Weaker expression is present in several muscles, including the flexor digitorum profundus (4), extensor metacarpi ulnaris (5), flexor carpi ulnaris (6), and triceps brachii (7). Strong expression is also observed in the integument surrounding the forelimb (8).

## Discussion

The present study revealed a broader *OSX/SP7* expression pattern during chicken embryogenesis than might be expected based solely on its established role in osteogenesis. Although *OSX/SP7* expression was observed in developing skeletal elements, consistent with its conserved function in bone formation, transcripts were also detected in several non-skeletal tissues, including the neuroepithelium, retina, central nervous system, muscles and integument. These findings suggest that the developmental roles of OSX/SP7 in birds may extend beyond the regulation of skeletogenesis.

The detection of *OSX/SP7* expression in developing skeletal elements is consistent with its well-established role in osteoblast differentiation and bone formation across vertebrates (Baek et al., 2010; Huang and Olsen, 2015; Nakashima et al., 2002; Niu et al., 2017; Renn and Winkler, 2014; Yu et al., 2017). This concordance validates both the specificity of the probe and the overall experimental approach, providing confidence that the broader expression domains observed in the present study reflect genuine aspects of *OSX/SP7* expression during chicken development rather than technical artefacts.

Particularly intriguing was the early expression of *OSX/SP7* within the developing neuroepithelium as the *OSX/SP7* transcripts were already detectable in the neural tube by embryonic day 2 and remained evident in selected regions of the central nervous system until at least HH40. Although the function of OSX/SP7 in neural tissues remains unclear, similar observations have been reported in other vertebrates. In zebrafish, *OSX/SP7* expression has been detected in the otic placode (DeLaurier et al., 2010), while in mice *OSX/SP7* expression has been described in the olfactory bulb, cortex and cerebellum up to at least 8 weeks of age (Park et al., 2011). In addition, RNA-sequencing data from human BioProjects (PRJNA280600 and PRJEB4337; see on https://www.ebi.ac.uk/ena/browser/view) indicate *OSX/SP7* expression within the central nervous system. Together, these observations suggest that neural expression of *OSX/SP7* may represent a conserved, albeit poorly understood, aspect of OSX/SP7 biology that extends beyond its classical role in osteogenesis.

One possibility is that this transient neural expression is associated with neural crest-related processes during early embryogenesis. However, lineage-tracing and functional studies will be required to determine whether these expression domains reflect direct roles in neural development or represent broader developmental regulatory programmes.

Interestingly, several members of the SP transcription factor family have been implicated in neural development and progenitor cell regulation, indicating that extra-skeletal functions within this family are biologically plausible (Bouwman and Philipsen, 2002; Göllner et al., 2001a; Göllner et al., 2001b; Johnson and Ghashghaei, 2020; Liang et al., 2013; Liu et al., 2018; Marin et al., 1997). Although the functional relevance of *OSX/SP7* expression in the developing nervous system remains unknown, the involvement of other SP family members in neural processes provides a broader biological context in which neural roles of OSX/SP7 merit further investigation.

Comparison of the present findings with reported *OSX/SP7* expression patterns in medaka and zebrafish reveals notable differences in the spatial and temporal regulation of *OSX/SP7* expression among vertebrates. In teleost fish, *OSX/SP7* expression first becomes apparent at the otic vesicle stage, corresponding to stages 28-30 in medaka and 14-16 somites (∼16-18 hpf) in zebrafish, and subsequently localizes to ossifying elements by stages 33-35 (∼3-4 dpf) in medaka and 55-72 hpf (∼2.5-3 dpf) in zebrafish (DeLaurier et al., 2010; Renn and Winkler, 2014). In contrast, avian ossification extends throughout the latter two-thirds of embryonic development and continues after hatching (Fratini et al., 2013; Nakamura et al., 2019; Sawad et al.), which is consistent with the expression observed in skeletal structures undergoing ossification in the present study. Whether the broader expression domains identified in chicken embryos reflect lineage-specific differences in the regulation and function of *OSX/SP7* remains an open question.

*OSX/SP7* expression was also detected in several muscle groups during later stages of chick embryonic development. To our knowledge, comparable patterns of *OSX/SP7* expression in embryonic muscle tissues have not been reported in other vertebrate species. At present, the biological significance of this observation remains unclear. Whether these expression domains reflect previously unrecognized functions of OSX/SP7 or represent lineage-specific regulatory features of avian development will require further investigation.

Overall, the expression patterns described here indicate that the developmental expression profile of *OSX/SP7* in chicken embryos is broader than traditionally appreciated. While its association with skeletal development is conserved, the presence of *OSX/SP7* transcripts in neural, muscular and integumentary tissues suggests that the functions of this transcription factor during avian embryogenesis may not be restricted to osteogenesis alone.

Comparison of the experimentally isolated *OSX/SP7* sequence with recently available genomic resources enabled reconstruction of the complete chicken *OSX/SP7* coding sequence and the inference of its associated untranslated regions from genomic data. Importantly, the same coding sequence was identified independently in both maternal and paternal haplotype-resolved chicken genome assemblies, providing strong support for the reconstructed ORF. These findings not only expand the available molecular resources for avian OSX/SP7 research but also provide a robust framework for future studies addressing the regulation and function of this transcription factor in birds.

Interestingly, the exon organization of *OSX/SP7* has been described differently in the literature, with both two-exon and alternative multi-exon structures reported in mammals. In chicken, our reconstructed chicken *OSX/SP7* transcript is most consistent with a two-exon organization, similar to that originally reported for human *OSX/SP7* by Gao et al. (Gao et al., 2004). Although the present findings do not resolve the broader question of *OSX/SP7* transcript organization across vertebrates, they suggest that this aspect of OSX/SP7 biology may be more nuanced than is generally appreciated.

The comparative analyses suggest that evolutionary constraints act differently across OSX/SP7 protein domains. Whereas the DNA-binding zinc-finger region remains highly conserved among vertebrates, the N-terminal portion of the protein appears considerably more divergent. Given that the N-terminal region has been proposed to participate in transcriptional regulatory functions and protein-protein interactions, this pattern of conservation may reflect differential selective pressures acting on distinct aspects of OSX/SP7 function during vertebrate evolution. However, the functional significance of these sequence differences remains to be established.

Taken together, our findings broaden the developmental landscape of *OSX/SP7* expression in birds and provide new molecular resources for future studies aimed at understanding the regulation and function of this transcription factor during vertebrate development.

## Author contribution

Conceptualization: C.W. A.W.; Investigation: D.L., C.Z., C.L., R.K., M.N., J. G., U.K.; Writing: C.L., M. N., C. W., J. G., C. Z., D. L., R. K., B. H., A. W.

## Animal ethics

This study was conducted in accordance with the German Animal Welfare Act (TierSchG) and the regulations of the Animal Care and Use Committee (ATV) of the local authorities (Regierungspräsidium Tübingen). Ethical approval was not required for this study under the regulations governing the use of early-stage avian embryos. Nevertheless, all procedures were carried out with due care and attention to animal welfare, and every effort was made to minimize potential suffering

